# High rates of evolution preceded shifts to sex-biased gene expression in *Leucadendron*, the most sexually dimorphic angiosperms

**DOI:** 10.1101/2021.01.12.426328

**Authors:** Mathias Scharmann, Anthony G Rebelo, John R Pannell

## Abstract

The males and females of many dioecious plants differ in morphological (Dawson and Geber 1999; Barrett and Hough 2013; Tonnabel et al. 2017), physiological (Juvany and Munné-Bosch 2015), life-history (Delph 1999), and defence traits (Cornelissen and Stiling 2005). Ultimately, such sexual dimorphism must largely be due to differential gene expression between the sexes (Ellegren and Parsch 2007), but little is known about how sex-biased genes are recruited and how their expression evolves over time. We measured gene expression in leaves of males and females of ten species sampled across the South African Cape genus *Leucadendron*, which shows repeated changes in sexual dimorphism and includes the most extreme differences between males and females in flowering plants (Midgley 2010; Barrett and Hough 2013; Tonnabel et al. 2014). Even in the most dimorphic species in our sample, fewer than 2% of genes showed sex-biased gene expression (SBGE) in vegetative tissue, with surprisingly little correspondence between SBGE and vegetative dimorphism across species. The identity of sex-biased genes in *Leucadendron* was highly species-specific, with a rapid turnover among species. In animals, sex-biased genes often evolve more quickly than unbiased genes in their sequences and expression levels (Ranz et al. 2003; Khaitovich et al. 2005; Ellegren and Parsch 2007; Voolstra et al. 2007; Harrison et al. 2015; Naqvi et al. 2019), consistent with hypotheses invoking rapid evolution due to sexual selection. Our phylogenetic analysis in *Leucadendron*, however, clearly indicates that sex-biased genes are recruited from a class of genes with ancestrally rapid rates of expression evolution, perhaps due to low evolutionary or pleiotropic constraints. Nevertheless, we also find evidence for adaptive evolution of expression levels once sex bias evolves. Thus, although the expression of sex-biased genes is ultimately responsive to selection, high rates of expression evolution might usually predate the evolution of sex bias.

## Introduction

Sexual dimorphism is common in dioecious plants, affecting a range of physiological (Juvany and Munné-Bosch 2015), morphological (Dawson and Geber 1999; Barrett and Hough 2013; Tonnabel et al. 2017), life-history (Delph 1999), and defence traits 6), yet most such cases of sexual dimorphism are less marked than they are in animals (Lloyd and Webb 1977; Moore and Pannell 2011; Barrett and Hough 2013). The South African genus *Leucadendron* stands out as an exception to this rule, with several of its approximately 80 dioecious species showing extreme levels of sexual dimorphism for leaf and plant architectural traits (Williams 1972; Rebelo 2001; Midgley 2010; Tonnabel et al. 2014) (Figure 1A). Previous work has found that species pollinated by wind tend to be more dimorphic than those pollinated by generalist insects (Tonnabel et al. 2014; Welsford et al. 2016), suggesting a possible role of intense male-male competition for siring success in selecting for divergence in morphology between males and females (Bond and Maze 1999). In *Leucadendron* species with specialised pollinators, male and female inflorescences may differ in attractive cues and provide different pollinator rewards (Hemborg and Bond 2005). Sexual dimorphism also tends to be greatest in species whose females maintain investment in woody seed cones for several years after they are produced (serotiny), suggesting that the relative cost of reproduction for males versus females has also played a role in divergence between the sexes (Harris and Pannell 2010).

**Figure 1.**
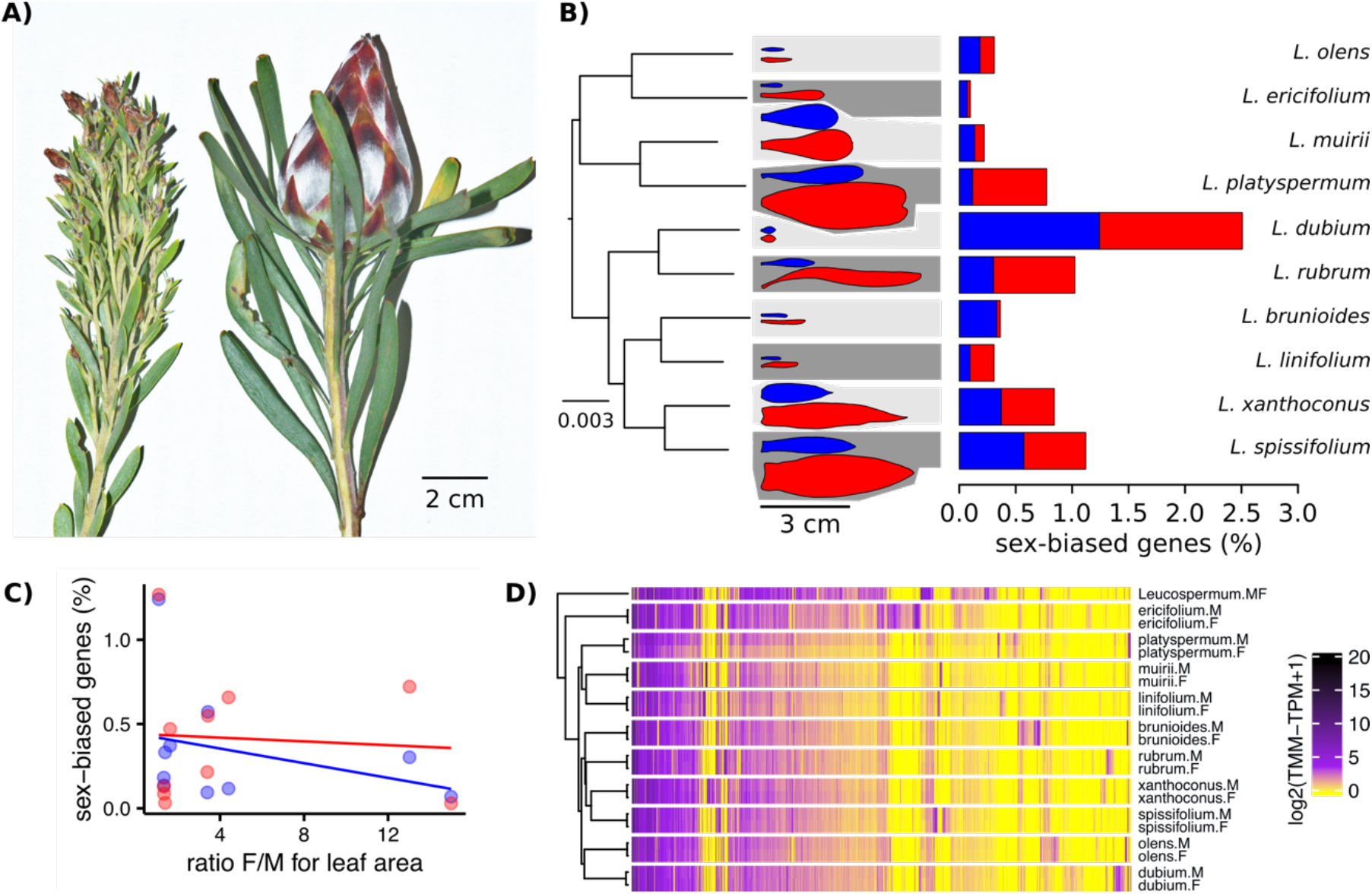
Extreme morphological sex-differences of the leaves evolved repeatedly in the genus *Leucadendron*, yet sex-biased expression affects only few genes, is not correlated with morphological dimorphism, and general male and female gene expression patterns do not exist. A). Typical male (left) and female (right) shoot tips of *L. rubrum*, a wind-pollinated species with extreme sexual dimorphism in leaves, stems and inflorescence. B) Left: super-matrix species tree; scale bar indicates expected number of substitutions per site. Middle: schematic examples of male (blue) and female (red) leaf outlines, to scale; background shading indicates lower sexual dimorphism as light grey and higher sexual dimorphism as dark grey. Right: Bar plots showing the percentage of male-biased (blue) and female-biased (red) genes among all expressed genes. C) The percentage of male-biased (blue) and female-biased (red) genes as a function of morphological sexual dimorphism, the ratio of female to male leaf area (Tonnabel et al. 2014). The lines show linear phylogenetic least squares fits for the percentage of female-biased transcripts in red (slope = −0.014, n.s.) and for the percentage of male-biased transcripts in blue (slope = −0.065, n.s.). D) Gene expression heatmap and hierarchical clustering dendrogram for the sexes of ten *Leucadendron* species and the hermaphroditic *Leucospermum* outgroup, for sex-biased genes only (650 genes). Gene expression values (columns) are the mean log_2_(TMM-TPM +1) per species and sex. The clustering of groups (rows) is based on distances calculated as 1 – Pearson’s correlation coefficient.

Morphological and physiological differences between the sexes of sexually dimorphic species must ultimately trace back to differences in gene expression (sex-biased gene expression, SBGE). Although some of these differences can be due to divergence between sex chromosomes (in certain species with genetic sex determination; (Bachtrog et al. 2014)), sex-biased genes tend to be largely autosomal (Ellegren and Parsch 2007; Mank 2009; Zemp et al. 2016). To the extent that morphological sexual dimorphism is related to SBGE, we might expect that species showing particularly striking differences between male and female in morphology would also be characterised by high levels of SBGE. To our knowledge, this prediction has hitherto been tested at a phylogenetic scale only for birds. Based on sample of six galloanserine species, Harrison et al. (Harrison et al. 2015) reported greater levels of SBGE in species with more exaggerated male traits, consistent with the hypothesis that the evolution of SBGE is driven by sexual selection. Whether such associations apply more generally is not yet known.

Sex-specific selection is thought of as the main evolutionary driver of sexual dimorphism (Barrett and Hough 2013), and the bird data (Harrison et al. 2015) appear to support the view that SBGE is similarly the target of some form of sexual selection. Comparisons of DNA sequences and gene expression phenotypes between species with contrasting degrees of sexual dimorphism can provide important evidence to evaluate this theory, because past phases of selection or drift can be diagnosed by characteristic patterns (Ranz et al. 2003; Khaitovich et al. 2005; Ellegren and Parsch 2007; Voolstra et al. 2007; Harrison et al. 2015; Naqvi et al. 2019). Under the hypotheses that sex-related adaptation drives SBGE, one would expect that sex-biased genes show elevated rates of amino acid substitution and faster rates of expression divergence between species (Pointer et al. 2013; Hollis et al. 2014; Immonen et al. 2014; Grath and Parsch 2016; Simmons et al. 2020). Tests of such predictions have provided evidence consistent with this idea in many animals (Ellegren and Parsch 2007; Harrison et al. 2015; Grath and Parsch 2016), but results have been mixed or negative in plants (Ellegren and Parsch 2007; Muyle 2019). Although an observation of rapid evolution of gene expression for sex-biased genes is consistent with the hypotheses of sexual selection, it remains unclear whether the high evolutionary rates found for sex-biased genes evolved prior to or after the evolution of sex bias (Orr 2000; Papakostas et al. 2014). This is an important uncertainty, because only acceleration of evolutionary rates that postdate or coincide with the evolution of sex-biased expression would be fully consistent with hypotheses invoking sex-specific (or sexual) selection) and thus the role of adaptation in the evolution of SBGE.

Here, we sequenced and analysed the transcriptome in the fully developed leaves of males and females of ten *Leucadendron* species sampled from across the genus to address two principle questions. First, we asked whether among-species variation in morphological sexual dimorphism corresponds to variation in levels of gene expression, as was found for birds (Harrison et al. 2015). Second, we inferred the history of recruitment, evolution and rate of turnover of genes with sex-biased expression across the genus to ask whether sex-biased genes acquired their characteristic rates of evolution prior to or after they became dimorphic in expression levels. Our study represents the first phylogenetic analysis of the evolution of sex-biased gene expression for plants, and our results urge caution in invoking adaptive evolution as the sole reason for high rates of expression evolution observed for sex-biased genes.

## Results and Discussion

We based our analysis on transcriptomes sequenced from species sampled from all major clades of the genus *Leucadendron*, recapturing variation generated over several tens of millions of years of evolution during the genus’ radiation (Sauquet et al. 2009) (reflected in up to 3.7% sequence differences at synonymous sites between species). We specifically paired closely related species that differed strongly in their level of sexual dimorphism. We also included in our sample one individual of the hermaphroditic species *Leucospermum reflexum* as an outgroup (sequence difference to *Leucadendron* spp. c. 5.3%).

### *Extent of sex-biased gene expression in* Leucadendron

De novo assembly of transcriptomes resulted in 73,307 to 570,280 contigs per species, of which 34,901 to 92,840 (13 – 48%) were identified as plant transcripts (Table S1). The abundant and diverse non-plant transcripts mainly belonged to fungi and bacteria, which is not surprising for wild, openly growing plants. We inferred and retained after filtering a set of 16,194 ‘orthogroups’ and quantified their expression across the eleven plant transcriptomes. We subsequently refer to these orthogroups as ‘genes’, though it should be noted that they are best understood as gene families rather than single genes. In total, we identified 650 genes that showed SBGE (Table S2), ranging from a minimum of seven male-biased and three female-biased genes in *L. ericifolium* (0.1% of expressed genes) to 138 male-biased and 141 female-biased genes in *L. dubium* (2.5%; Figure 1B). Similar patterns were revealed when sex-biased genes were counted, or when the magnitude of bias (fold-changes) was cumulated over the genes (Table S3).

Although *Leucadendron* represents some of the most extreme examples of morphological sexual dimorphism recorded in flowering plants (Bond and Maze 1999; Harris and Pannell 2010; Midgley 2010; Barrett and Hough 2013), our results indicate that the genus is not at all exceptional in terms of the extent of SBGE within species, including the most morphologically dimorphic ones. For instance, our results compare with the low estimates for *Populus* (Robinson et al. 2014) and *Salix* (Darolti et al. 2018; Sanderson et al. 2018), in which < 0.1% of genes are sex-biased, as well as the somewhat higher estimates reported for *Silene latifolia* (Zemp et al. 2016) and *Mercurialis annua* (Cossard et al. 2019), which have about 2% of sex-biased genes. In contrast with these dioecious plants, even non-reproductive tissues of animals with separate sexes can express hundreds or thousands of genes with a sex-biased pattern (Naqvi et al. 2019).

### Correspondence in sexual dimorphism between morphology and gene expression

Previous work on birds has demonstrated a positive correspondence between variation in SBGE and levels of sexual dimorphism for a number of traits related to the inferred intensity of sexual selection (Harrison et al. 2015). To test whether this finding might also apply to a plant genus with extreme among-species variation in levels of sexual dimorphism, we asked whether convergent evolution of sexual dimorphism in *Leucadendron* for morphology is associated with convergence of SBGE. Our data clearly reject this hypothesis: morphological sexual dimorphism, measured either as the ratio of female to male leaf area, or female to male specific leaf area (leaf mass per area), were correlated neither with the number, proportion nor the cumulative foldchanges of sex-biased genes (phylogenetic least-squares regressions, all *P*> 0.05, Figure 1C, Figure S1). For instance, *L. ericifolium* shows extreme morphological dimorphism but had only seven male-biased and three female-biased genes (0.1% of expressed genes), whereas *L. dubium* shows very little evidence for morphological dimorphism but had 138 male-biased and 141 female-biased genes (2.5%). We also failed to find any genes that were consistently sex-biased in species with either low or high morphological dimorphism; instead, the identity of sex-biased genes was largely unique in each species. Annotation of *Leucadendron* sex-biased genes (SBGs) against *Arabidopsis thaliana* genes revealed very diverse putative functions (Tables S2, S5, S6). While there were some obvious sex-differences within species, such as male-bias for phenylpropanoid / flavonoid biosynthesis and female-bias for carbon assimilation in *L. dubium* (Table S7), there were no clear male or female functional patterns across species (Figure S2). On the contrary, the two gene functional clusters most strongly overrepresented in sex-biased genes as compared to all genes were the same among the male- and female-biased genes over all ten species (involving flavonoid biosynthesis and secreted extracellular proteins; Tables S5 and S6). This hints at the possibility that molecular sexual dimorphism could to some degree be reversed between different species. While our analysis is based on leaf traits, we conjecture that SBGE is also unlikely to be related to dimorphism for other traits, such as height and branching architecture, because dimorphism in these traits tends to be correlated with leaf sexual dimorphism (Bond and Midgley 1988; Harris and Pannell 2010).

### *Evolution of SBGE across the genus* Leucadendron

To gain further insight into the evolution of SBGE in the *Leucadendron* radiation, we clustered species and sexes according to gross similarity in gene expression. The males and females of a given species were more similar in their expression profiles than were individuals of the same sex of different species, i.e., clustering grouped samples by species rather than by sex (Figure 1D). At the level of species, clustering based on gene expression failed to reflect the DNA sequence phylogeny, indicating relatively little phylogenetic inertia in expression levels. Overall, there were no clear general male-like or female-like gene expression profiles shared across the genus, and the convergence towards sexual dimorphism observed at the morphological level is not apparent at the level of gene expression. This result parallels the low number of conserved SBGE in animals for non-reproductive tissues (Harrison et al. 2015; Naqvi et al. 2019). In contrast, gene expression in bird gonads has been found to cluster by sex rather than by species, consistent with evolutionary conserved differences for genes expressed in testes and ovaries (Harrison et al. 2015).

Our genus-wide sampling allowed us to consider the ancestral versus derived nature of SBGE for the sequenced genes, using maximum likelihood to reconstruct ancestral expression states for each sex-biased gene (Table 1, Figure S3). Of the 650 genes that showed SBGE in at least one species, only 63 (9.7%) were in two or more species, and in only four out of these 63 sex-biased genes was the bias likely to have been inherited from a common ancestor. Furthermore, our analysis suggests that none of the sex-biased genes we detected across any species has been sex-biased in the common ancestor of the genus (Figure S3), despite the fact that dioecy was probably the ancestral state for *Leucadendron*. Note that this result does not imply that ancestral *Leucadendron* had little SBGE, because our reconstruction is necessarily limited to recent SBGs (Harrison et al. 2015).

**Table 1.** Summary of inferred evolutionary histories for 650 sex-biased genes in *Leucadendron*. Permutations were used to generate numbers of genes in each category expected under the null hypothesis that the identity of SBGs is random within each species, excluding the three genes that gained sex-bias only ancestrally. All observed counts were significantly different from the null. The phylogenetic patterns of shared and divergent SBGE are shown in Figure S3.

| category of sex-bias | observed | mean of expected | p-value (one-sided) |
| --- | --- | --- | --- |
| uniquely male-biased, gained at tip | 265 | 312 | $< 1 \times 10^{-5}$ |
| uniquely female-biased, gained at tip | 322 | 377.5 | $< 1 \times 10^{-5}$ |
| shared male-biased, single ancestral gain | 3 | NA | NA |
| shared female-biased, single ancestral gain | 1 | NA | NA |
| shared male-biased, repeatedly gained (ancestral and/or tip) | 12 | 2.4 | $< 1 \times 10^{-5}$ |
| shared female-biased, repeatedly gained (ancestral and/or tip) | 19 | 3.7 | $< 1 \times 10^{-5}$ |
| divergent sex-biased (gains ancestral and/or tip) | 28 | 6.1 | $< 1 \times 10^{-5}$ |

Although our results indicate that SBGE largely evolved in an idiosyncratic, lineage-specific manner in *Leucadendron*, certain genes were more likely to evolve SBGE than others. Fifty-nine out of the 63 genes showing sex-bias in more than one species acquired sex-bias more than once (Table 1, Figure S3, Table S4). About half of these (31 genes) acquired sex-bias of the same direction in distinct species. While convergent sex-bias was limited to two species in most genes, one component of the respiratory chain (OG0010406, corresponding to *Arabidopsis thaliana* gene ATMG00513) evolved female-bias in three species, and another gene, putatively involved in phosphate homeostasis, evolved female-bias independently in four species (OG0000761, SPX3, AT2G45130). Phosphorous metabolism is known to be sexually dimorphic in other dioecious plants (Zhang et al. 2014; Zhang et al. 2019). Interestingly, the remaining 28 genes with apparent convergence in SBGE actually showed reversed sex-bias, i.e. they evolved to male-bias in some species but to female-bias in others. While most of these genes had sex-reversed expression in only one pair of species, six genes were sex-biased in opposite directions in three species (i.e. twice female-biased and once male-biased). Four of these genes are involved in flavonoid biosynthesis (OG0009936, OG0001220, OG0001123, OG0004450; corresponding to AT4G22880, AT5G42800, AT1G08250, AT1G61720), and thus have putative functions attributable to leaf pigmentation, which is sexually dimorphic in several *Leucadendron* species (Rebelo 2001). The other two genes with reversed sex-bias in three species are putatively involved in cytokinin biosynthesis (OG0001906; AT4G35190), and a vacuolar sucrose invertase (OG0000569, AT1G62660). The extent of interspecific overlap in similarly sex-biased and reversed sex-biased expression in terms of gene number was substantially and statistically significantly greater than expected by chance; correspondingly, the number of uniquely SBGs was lower than expected (permutation tests, all *P*< 10^-5^; Table 1). It is thus clear that the set of genes recruited for SBGE during the radiation is much narrower than it could have been.

### *Origin and evolutionary rates of sex-biased genes in* Leucadendron

Although dioecy is likely the ancestral sexual system in *Leucadendron* (Sauquet et al. 2009; Tonnabel et al. 2014), and strong sexual dimorphism has evolved several times independently across the genus, our results clearly indicate that sex bias in the expression of individual genes is neither ancestral nor highly convergent. This pattern points to a high rate of turnover of sex-biased expression among genes. In animals, sex-biased genes often show more rapid sequence evolution than un-biased genes, which could reflect positive selection or relaxed purifying selection (Ellegren and Parsch 2007; Naqvi et al. 2019). In contrast, plants have so far not revealed a similar tendency (Zemp et al. 2016; Cossard et al. 2019; Muyle 2019). Consistent with these previous findings for plants, and again in contrast to animals, we too found no evidence for faster sequence evolution of sex-biased genes in *Leucadendron* (Figure S4). It is possible that sex-biased genes do evolve faster in flowering plants, but dioecy in plant is typically an evolutionarily recent trait (Renner 2014; Charlesworth 2016), making detection of faster evolution difficult. In *Leucadendron*, in which dioecy might be several tens of million years old (Sauquet et al. 2009; Tonnabel et al. 2014), our results indicate that SBGE is much more recent than the evolution of separate sexes at the base of the genus.

In stark contrast to the lack of evidence for faster sequence evolution of sex-biased genes in *Leucadendron*, we found that they have in fact been evolving more quickly than unbiased genes in terms of their expression levels (Figure 2). More rapid evolution of expression for sex-biased genes, too, is common in animals. This has been documented for whole-body transcriptomes of *Drosophila* (Ranz et al. 2003), as well as vertebrates, in which testes gene expression diverges faster between species than expression in non-reproductive tissues (Khaitovich et al. 2005; Voolstra et al. 2007; Harrison et al. 2015). Less is known about the rates of expression evolution for sex-biased genes in plants, but a comparison between two species of *Silene* indicated that their rates are higher than those of unbiased genes (Zemp et al. 2016). Our study across a diverse clade of dioecious plants adds substantially to the modest sampling in plants so far and suggests that expression evolution is likely more rapid for sex-biased genes quite generally, and that it evolves more quickly than the gene sequences themselves.

**Figure 2.**
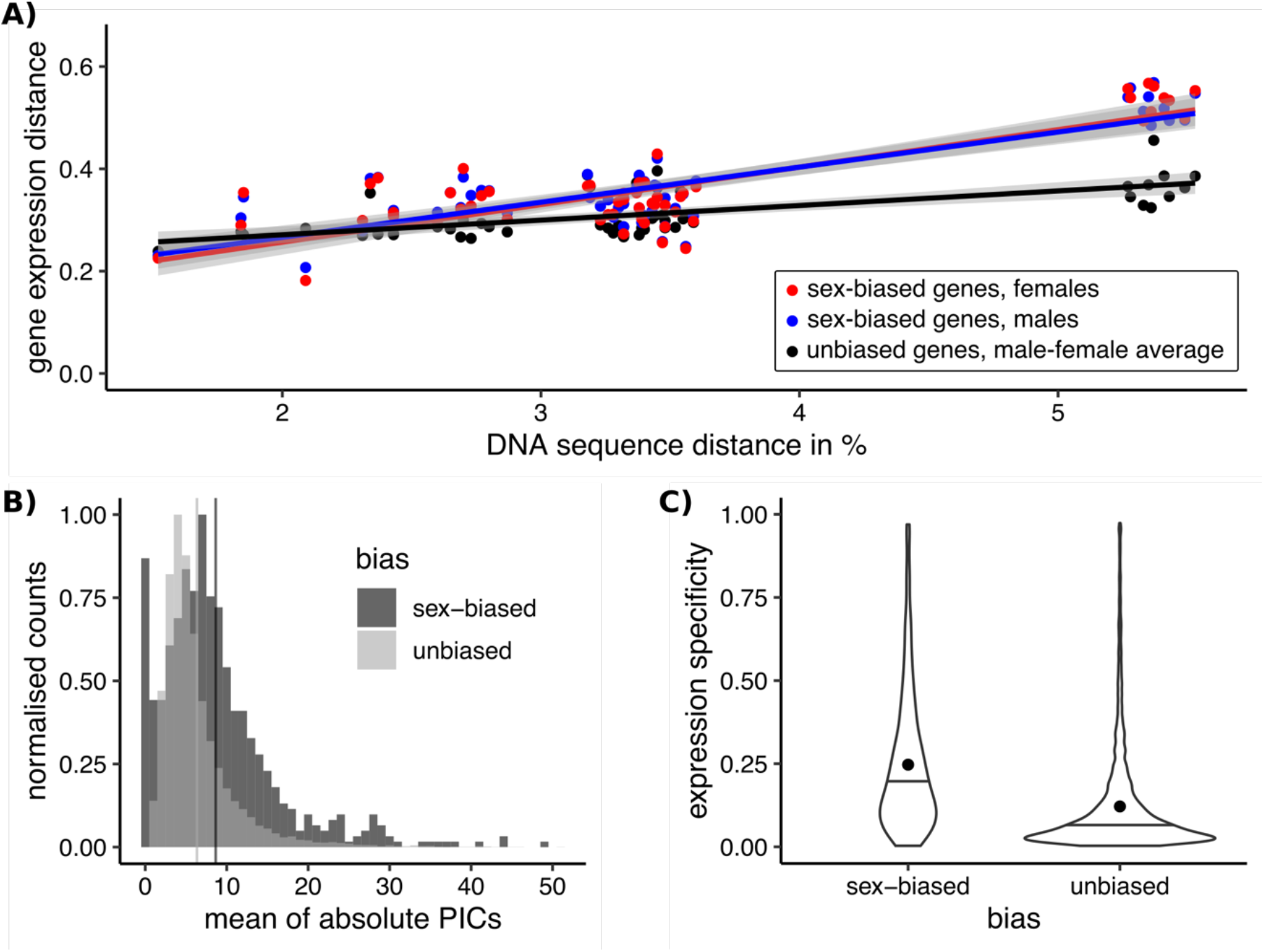
Sex-biased genes in *Leucadendron* have ancestrally and intrinsically higher rates of expression evolution and higher specificity of expression. A) Interspecific gene expression distance as a function of DNA sequence distance for 55 species pairs of dioecious Leucadendron and the hermaphrodite relative *Leucospermum*. Sex-biased expressions themselves were excluded when calculating gene expression distances. Gene expression distances are shown for three different categories: distance between species (mean over the sexes) for unbiased genes, distance between males for sex-biased genes, and distance between females for sex-biased genes. DNA distances > 4% pertain to *Leucadendron-Leucospermum* pairs. The shaded envelopes around linear regression lines represent parameter standard errors. DNA sequence distance was significantly correlated with each of the three categories of gene expression distance (Mantel tests, all p <= 0.0225). B) Histograms of mean absolute standardized phylogenetically independent contrasts (PICs) of gene expression for sex-biased and unbiased genes. The difference in means of the two categories is 2.3 (permutation p = 2 x 10^-5^). C) Expression specificity (Shannon entropy) over different tissues and developmental stages for sex-biased and unbiased genes in Leucadendron. Violins show the complete distributions, horizontal bars indicate the medians and dots the means. The difference of the means is 0.095 (permutation p = 2 x 10^-5^).

In contrast to previous work in plants and animals, the phylogenetic context of our sampling allowed us to ask whether rates of expression evolution accelerated for genes once they became sex-biased, or whether expression levels were already evolving rapidly before they became sex-biased. The former possibility would be consistent with the outcome of sex-specific (or sexual) selection, for example (Pointer et al. 2013; Hollis et al. 2014; Immonen et al. 2014; Simmons et al. 2020), whereas the latter possibility would be more consistent with reduced functional constraints of expression levels for sex-biased genes, i.e., they evolve more quickly because they can do so with impunity (Orr 2000; Papakostas et al. 2014), and hence then are freer to become sex-biased, too. We thus compared rates of expression evolution between unbiased genes and SBGs. Importantly, we here inferred the rates for SBGs only among species in which these genes were actually unbiased, an analysis made possible by the largely species-specific nature of sexbias in *Leucadendron*. Phylogenetically independent contrasts (Garland 1992) revealed that rates of expression evolution in *Leucadendron* tend to be higher for sex-biased than unbiased genes in general (Figure 2 B). Moreover, the interspecific expression distance of sex-biased genes was generally greater, and increased more steeply with interspecific sequence divergence (i.e., with evolutionary time; Figure 2 A). Because these rates of expression evolution for SBGs were inferred only from expression values in species not showing sex-bias, it is clear that the elevated rates preceded these genes’ acquisition of sex-bias, and are independent of it.

Rates of expression evolution inferred from differences among species might appear higher for those genes that show greater within-species variation in expression; indeed, intra-group variation in expression level generally correlates positively with inter-group differences (Figure S5, see also (Uebbing et al. 2016)). In our study, variation in expression among the males or females for a given species of *Leucadendron* was sometimes greater, similar or lower in sex-biased genes than for unbiased genes (Figure S6). Thus, we tested whether the slope of inter-specifc expression distance against sequence divergence in SBGs was greater than slopes of random subsets of unbiased genes that were matched to the SBGs in their degree of intra-sex expression variation. We found that all slopes of 1,000 of such random but noise-matched gene subsets were lower than the slopes of the actual SBGs (Figure S7). This result implies that the observed faster rate of expression evolution in *Leucadendron* is a real distinguishing feature of its SBGs and not merely an artefact of their expression noise. Moreover, permutations verified that there were more sex-biased genes (in most species; Figure S8) than expected by chance alone. Overall, we are thus confident that the inferred higher ancestral rates of expression evolution for sex-biased genes are real and not an artefact of analysis or of sampling genes with especially large expression variation.

Sex-biased genes in animals are frequently associated with higher expression specificity across a range of tissues and developmental stages (Orr 2000; Papakostas et al. 2014), suggesting that their higher evolutionary rates may be due to reduced pleiotropic constraint (Mank and Ellegren 2009; Parsch and Ellegren 2013; Naqvi et al. 2019). We thus asked whether the sex-biased genes in *Leucadendron* had similarly greater expression specificity, using tissue specificity in the hermaphrodite *Arabidopsis thaliana* (Klepikova et al. 2016) and sequence similarity to *Leucadendron* to roughly estimate ancestral tissue specificity in *Leucadendron*. With this proxy, the ancestral expression specificity of sex-biased in *Leucadendron* was indeed greater than that for unbiased genes (Figure 2 C). This result is consistent with the possibility that sex-biased genes in *Leucadendron* are pleiotropically less constrained in their expression levels than unbiased genes. Because all cases of sex bias in *Leucadendron* are recently evolved, we infer that the lower pleiotropic constraint to which these genes are subject must have preceded the acquisition of sex bias, rather than being a consequence of sex-biased expression.

If sex-biased expression evolved in genes for which expression levels were already evolving quickly, we must consider that sex bias is the result of neutral evolution in gene expression (Nourmohammad et al. 2017), e.g., due to relaxed constraint. To further investigate the hypothesis that sex-biased expression had evolved neutrally in *Leucadendron*, we compared shifts in expression that led to sex-biased expression (sex-biased shifts) with those that did not (unbiased shifts). We defined an evolutionary shift in expression as a difference of at least 50% between the mean expression values of sister species. Note that sex-bias is here narrowly defined for the species under scrutiny, i.e., a given gene could be classed as undergoing a sex-biased shift for one species but unbiased shifts for the nine others, or not undergoing any shift at all. We calculated for each expression shift the delta-x statistic (Hsieh et al. 2003; Rifkin et al. 2003; Moghadam et al. 2012) for an indication of whether it might have been driven by natural selection. Reflecting the notion that directional selection should both increase divergence between groups and decrease variation in the group under selection, delta-x takes higher values the higher divergence and the lower polymorphism. Low expression polymorphism relative to interspecific divergence is a hallmark of adaptive evolution (Nuzhdin et al. 2004; Ometto et al. 2011; Khodursky et al. 2020).

We found that putatively adaptive expression shifts leading to sex-bias occurred in both males and females. In particular, c. 21% of shifts leading to SBGE were putatively adaptive in at least one of the sexes, choosing the threshold of divergence five times higher than polymorphism (delta-x = 5.0) to class expression shifts as adaptive (Table 2). However, the incidence of signatures consistent with adaptation in expression levels was significantly higher among shifts not leading to sex-bias (Table 2). This result is extremely robust to the choice of threshold for classing expression differences between species as shifts, and for classing shifts as adaptive. The pattern soundly rejects the hypothesis that sex-biased expression in *Leucadendron* is generally a result of neutral evolution. Instead, it would appear that while adaptations that establish sex-biased expression did occur, shifts towards sex-biased expression were less frequently adaptive than expression shifts unrelated to gender or sexual dimorphism.

**Table 2.** Counts and chi-squared tests for shifts in gene expression in ten *Leucadendron* spp. as classified by an index of selection and by sex-bias. Shifts are defined as differences of 1.5-fold or greater in mean expression between sister species. Delta-x, an index of selection, is the ratio of absolute difference in expression between sister species to the standard deviation of expression in a focal species. Shifts in the category ‘high delta-x’ (here set to 5.0 and greater) are more consistent with directional selection (adaptation) than ‘low delta-x’ shifts.

|  |  | Male expression shifts |  | Female expression shifts |  |
| --- | --- | --- | --- | --- | --- |
|  |  | sex-biased | unbiased | sex-biased | unbiased |
| observed | high delta-x | 103 | 22349 | 100 | 21639 |
|  | low delta-x | 394 | 34002 | 373 | 36412 |
| expected | high delta-x | 196.3 | 22255.7 | 175.7 | 21563.3 |
|  | low delta-x | 300.7 | 34095.3 | 297.3 | 36487.7 |
| chi-square Yates, df = 1 |  | 73.134 |  | 51.622 |  |
| p |  | < 2.2 x 10<sup>-16</sup> |  | 6.73 x 10<sup>-13</sup> |  |

### Concluding remarks

Our study represents the first genus-wide comparative study of the evolution of SBGE in plants. In common with the findings of previous studies of one or two species, our results confirm that SBGE in vegetative tissues is not as pronounced as it sometimes is in animals. We failed to find correlations between SBGE and morphological dimorphism, perhaps because the small and large leaves of *Leucadendron* males and females mostly differ in cell number but not cell type composition and cell physiology, unlike the gonads of animals in which such correlations exist (Harrison et al. 2015; Montgomery and Mank 2016). We also failed to find evidence for more rapid sequence evolution of sex-biased genes compared with unbiased genes, in contrast with observations for sex-biased genes in animals. By taking a phylogenetic perspective however, we were able to consider shifts in gene expression over evolutionary time in independent lineages with common ancestors, leading to several novel insights. These include an almost complete lack of convergent evolution of sex-bias of individual genes, despite striking convergence in aspects of morphological dimorphism across the genus, and that sex-biased genes are recruited from a class of genes with special intrinsic and ancestral properties.

Most significantly, our analysis indicates that the rapid rates of expression evolution for sex-biased genes generally predate their recent acquisition of sex bias itself, suggesting that sex bias may evolve more easily in genes that are intrinsically less constrained in relative expression level. The hypothesis of reduced constraint, which is also supported by the fact that the sex-biased genes identified in *Leucadendron* tend to have greater expression specificity over various plant tissues and developmental stages than unbiased genes, suggests that much of the SBGE observed in plants (and perhaps animals too) may in fact have only limited functional importance. On the other hand, gene expression divergence over polymorphism ratios suggested that at least some of the expression shifts to sex-bias were adaptive in either one or both sexes. Thus, while drift in expression likely contributes to variation in gene expression and could explain much of the striking patterns of turnover among sex-biased genes we have identified in *Leucadendron*, the potentially random walks in expression space of these genes have not, however, been entirely invisible to the eye of sex-specific selection.

## Materials and Methods

Mature leaves of *Leucadendron* were sampled in South Africa, and kept for a brief period at 0°C in nucleic acid preservation buffer before storage at −80°C. *Leucospermum* was sampled in the Botanical Garden of Zurich. We constructed 120 stranded RNA-seq libraries from total RNA, which were sequenced for 150 bp paired-end reads on six lanes of an Illumina HiSeq4000. Data are deposited under the ENA project TBA.

De-novo transcriptomes were assembled for each species separately using Trinity (Haas et al. 2013). Contigs were classifed as contaminant micro-organisms or else plant-derived by BLAST search against the NCBI nt database. Orthogroups, i.e. genes in the sense of this study, were inferred from protein-coding plant contigs with OrthoFinder (Emms and Kelly 2019).

Gene expression was quantified (Patro et al. 2017) using the species-specific de-novo transcriptomes, converted to read counts (‘scaledTPM’), and summarized to gene level, i.e. the orthogroups. Thus, the expression level of a gene in this study is a weighted summary of the expression over all contigs belonging to an orthogroup.

Sex-bias was tested in each species separately using TMM-normalised (Robinson et al. 2010) ‘scaledTPM’ read counts, by differential gene expression analysis in edgeR (Robinson et al. 2010) with the exactTest function, for n = 6 per sex for *L. rubrum*, *L. spissifolium*, *L. olens, L. brunioides, L. linifolium, L. muirii, L.dubium* and *L. xanthoconus*, and six males versus five females in *L. platyspermum* and *L. ericifolium*. Sex-bias was considered significant at a false discovery rate (FDR) of 5% (Benjamini and Hochberg 1995) and a minimum two-fold change between the sexes.

For further analyses, the expression levels of 16,194 genes were normalised for gene length, library size and compositionality over all samples of all species together, resulting in TMM-normalised TPM values, and transformed by log_2_(x+1). ComplexHeatmap (Gu et al. 2016) was used to visualise gene expression and to cluster species and sexes by gene expression distance. The transcriptomes contained 3,032 one-to-one orthologs, which we used to infer a concatenation species tree by maximum-likelihood, and to count DNA sequence divergence at four-fold degenerate sites between all pairs of taxa.

We regressed leaf sexual dimorphism of *Leucadendron* species (morphological / macroscopic dimorphism), here the ratio of female to male leaf size (Tonnabel et al. 2014), or specific leaf area, against the number of male-biased and female-biased genes using a phylogenetic least squares model (Pinheiro et al. 2018).

To infer where along the species tree genes had evolved sex-bias we performed categorical ancestral trait reconstruction (Paradis and Schliep 2019). To test whether the observed frequencies and gene identities of repeated and divergent evolution of sex-bias were different from a random expectation, we devised a permutation procedure.

Functional annotation of *Leucadendron* genes was performed against *Arabidopsis thaliana* genes, and a broad level functional categorization was generated through the TAIR web portal and the DAVID database (Huang et al. 2009), which was also used to cluster gene functional annotations.

Rates of gene expression evolution were quantified by two measures: First, we calculated the mean of the absolute standardized phylogenetically independent contrasts (PICs, (Garland 1992)) per gene, and differences between sex-biased and unbiased genes were tested by permutation. Second, gene expression distances between pairs of species were calculated as 1 – Pearson’s correlation coefficient between the expression values. This was done for three different categories of genes, namely for unbiased genes with the average expression levels over both sexes, for sex-biased genes with the male expression levels, and with the female expression levels. Both expression evolution rate measures for sex-biased genes were based only expression levels and species in which these genes were not sex-biased; hence sex-biased expression itself did not contribute to a gene’s expression evolutionary rate estimate.

Correlations between DNA sequence and gene expression distance were visualised by linear model fits, and significance evaluated by Mantel tests. To verify that higher expression evolutionary rates observed for sex-biased genes were not spurious artefacts, we devised a series of sanity checks.

As an indicator of directional selection on gene expression level, we used delta-x (Zemp et al. 2016), here the ratio of the difference in mean expression between two groups (divergence) over the standard deviation within a focal group (diversity / polymorphism). Higher values of delta-x are more consistent with adaptive evolution.

The expression specificity for *Leucadendron* genes was transferred from *Arabidopsis thaliana* genes, as calculated over the Klepikova expression atlas (Klepikova et al. 2016). Differences in expression specificity between sex-biased and unbiased genes were assessed by permutation. Rates of coding sequence evolution for sex-biased and unbiased genes were estimated using PAML (Yang 2007) and compared between the two classes of genes by permutation.

A detailed description of the methods of this study is provided in the Supplementary Information.

## Acknowledgments

We thank Yves Cuenot and Dessislava Savova-Bianchi for assistance in the lab, and members of the Pannell lab, especially Jeanne Tonnabel, for advice and helpful discussions. We are grateful to Jörn Gerchen, Nora Villamil Buenrostro, Xinji Li, Michael Lenhard, Darren Parker and Guillaume Cossard for helpful comments on an earlier version of the manuscript. Sequencing was performed by the Functional Genomics Center Zurich (FGCZ) and bioinformatic analysis was carried out on the servers of the Division Calcul et Soutien à la Recherche (DCSR) at the University of Lausanne. The research was funded by a grant to JRP by the Swiss National Science Foundation (SNF grant number 310030_185196).

## Author Contributions

JRP and MS conceived the study and wrote the manuscript. MS conducted wet lab procedures, bioinformatics and statistical analyses. All authors conducted the field work. AGR contributed extensive knowledge about the sampling localities and ecology of the species sampled. All authors read and contributed to the final stages of manuscript preparation.

## Supplementary Information

### SI Text Materials and Methods

#### Plant material and RNA-seq

Based on a phylogenetic tree and analyses of trait evolution in *Leucadendron* (Tonnabel et al. 2014), we selected five phylogenetic pairs of species displaying low and high leaf-size dimorphism. Leaf material from these ten species was collected from wild populations in their natural habitats in South Africa, with males and females sampled in an alternating fashion along transects through the respective population. For an outgroup, we sampled leaf material from a single individual of the hermaphroditic species *Leucospermum reflexum* from the Botanical Garden of the University of Zurich, Switzerland.

For all species, we removed mature leaves just below the most recent inflorescences, cut them into pieces of < 5 mm length, and immediately submerged the material in 8 mL tubes (Sarstedt 60.542.007) in ice-cold RNA-later (Ambion) or a homemade nucleic acid preservation buffer (Camacho-Sanchez et al. 2013). We used a manual vacuum pump to enhance infiltration of the leaves by the buffer. The samples from one population were all collected over a period of approximately two hours. Sample tubes were kept at 0°C for up to 6 days and then frozen at −80°C until further use. Total RNA was extracted from the pickled frozen leaves by grinding to a fine powder under liquid N2 and purified with the Maxwell 16 LEV Plant RNA Kit (Promega Corporation, Madison, WI USA) using a KingFisher Duo Prime robot (Thermo Fisher Scientific, Waltham, MA USA). The RNA extracts showed generally high integrity (Bioanalyzer profiles). Sequencing libraries were constructed using the KAPA Stranded RNA-Seq kit (KAPA Biosystems, Wilmington, MA USA) with Illumina-compatible indexed Pentadapters (PentaBase ApS, Odense, Denmark). A total of 120 libraries were multiplexed in six pools of 20 each, such that each pool contained one male and one female from each species, to avoid batch effects. Each pool was sequenced in a separate lane for 150 bp paired-end reads on an Illumina HiSeq 4000 at the FGCZ.

#### Transcriptome de-novo assemblies and identification of contaminant sequences

Adapter sequences were trimmed from the raw reads by trimmomatic (Bolger et al. 2014), and transcriptomes were assembled de novo for each species separately with the data of three males and three females (*Leucospermum*: one individual), using Trinity 2.5.1 (Haas et al. 2013). To remove contamination from epi- and endophytic micro-organisms, all Trinity contigs were searched against NCBI nt (Jan 26, 2017) using NCBI BLAST 2.7.1 (Altschul et al. 1990), with an e-value threshold of 1 x 10^-5^, recording the taxonomy identifier of the best hit. Sequences were classified as ‘Viridiplantae’ or not, using NCBI taxonomy.

#### Inference of orthogroups

We used OrthoFinder v2.3.3 (Emms and Kelly 2019) to identify orthogroups, i.e. groups of homologous sequences descending from a common ancestral gene in our set of eleven species (i.e., including the *Leucospermum* outgroup). To this end, de novo contigs, assembled by Trinity (see above), were filtered for those classified as of Viridiplantae origin, and contig redundancy was reduced by selecting the longest isoform of each gene. On these, open reading frames (ORFs) were predicted using TransDecoder v5.5.0 (Haas et al. 2013), considering all ORFs >100 bp long, and shorter ORFs if they were supported by a homology search against plant proteins in SwissProt or ARAPORT11 (blastp, e-value cutoff 10^-5^). The set of predicted protein sequences from each of the eleven species was then supplied to OrthoFinder, which was run with default settings, and produced 26,553 raw orthogroups. We note that the number of Viridiplantae transcripts assembled by Trinity varied about threefold between species. This is likely mainly due to technical effects and differences in expression state of genes rather than differences in the gene content of the species’ genomes, because *Leucadendron* (2n=26 (Liu et al. 2006)) as well as *Leucospermum* (2n=24 (Rourke 1970)) are generally diploid.

#### Quantification of gene expression levels

Gene expression was quantified using pseudo-alignment with Salmon (Patro et al. 2017), and with the specific Trinity assembly for each species as a reference. The salmon option for metagenomic datasets (“--meta”) was used, because the references contained all contaminant contigs and (partially) redundant homologous sequences. This strategy allows for maximum mapping success and avoids mis-alignment of contaminant reads to plant contigs. Using the R package tximport (Soneson et al. 2016), we converted Salmon abundance estimates to read counts (“scaledTPM”), and summarized to gene level, i.e. the OrthoFinder orthogroups plus the two additional categories “non-coding plant transcript” and “contaminant”. Our strategy to use all sequences and then summarize the read counts with tximport does not distinguish among orthologs, paralogs, isoforms, splice variants or any other form of homology; the expression value of a ‘gene’ is thus a weighted summary of the expression counts for all sequences in that orthogroup. Another consequence of the tximport summarization within orthogroups is that eventual strong differences in only some of the homologs within an orthogroup are ‘diluted’ in the summarized count. This would constitute a conservative bias for inference of differences in expression.

#### Differential expression tests

Sex-bias was tested for each species separately by differential gene expression analysis in edgeR (Robinson et al. 2010), with n = 6 per sex for *L. rubrum*, *L. spissifolium*, *L. olens, L. brunioides, L. linifolium, L. muirii, L.dubium* and *L. xanthoconus*, and six males versus five females in *L. platyspermum* and *L. ericifolium*. The ‘scaledTPM’ read counts from tximport were used. Within species, only genes passing a minimum expression cut-off (the count per million corresponding to more than ten mapped reads in the smallest library) in at least three samples were considered as significantly expressed and included in the test; genes with zero in one of the sexes were thus automatically included. Due to this filtering, only 16,194 of the 26,553 raw orthogroups were tested for differential expression in at least one species. The data were TMM-normalised (Robinson et al. 2010) to account for compositionality as well as library size. Differential expression was tested with the exactTest function and tag-wise dispersion estimate. Sex-bias was considered significant at a false discovery rate (FDR) of 5% (Benjamini and Hochberg 1995) and a minimum two-fold change between the sexes.

#### Expression data preparation for analyses other than differential expression tests

For all downstream analyses other than differential expression tests, expression values were normalized for library size, gene lengths, and compositionality. In particular, we first filtered the genes to those included in the differential expression test of at least one species (see above: 16,194 genes), and then used edgeR to apply TMM normalization factors over the Salmon TPM values of all 119 samples together. The resulting TMM-normalised TPM values were transformed by log_2_(x + 1). We note that this strategy included genes that had zero expression in some species. ComplexHeatmap v2.2.0 (Gu et al. 2016) was used to visualise gene expression data, and to cluster species and sexes by gene expression distance (1 – Pearson’s r).

#### Species tree and interspecific sequence divergence

The orthogroups contained 3,032 one-to-one orthologs which were present in each of the eleven taxa. These ortholog sequences were cleaned from putatively non-homologous domains using PREQUAL v1.02 (Whelan et al. 2018), aligned with MAFFT v7.455 (Katoh and Standley 2013) on the peptide level, and finally back-translated to coding sequences. A species tree was estimated from a concatenated supermatrix using RAxML v8.2.12 (Stamatakis 2014) with the GTRCAT substitution model. Uncertainty was evaluated with RAxML’s SH-like algorithm. The same supermatrix was used to count raw sequence divergence at four-fold degenerate sites between all pairs of taxa.

#### Comparison of leaf dimorphism against sex-biased expression

Leaf size dimorphism was quantified as the ratio of female to male leaf size, using measurements taken from (Tonnabel et al. 2014). As a second measure of leaf dimorphism, we used the ratio of female to male specific leaf area (SLA, g/cm^2^). Several fresh or wet-preserved leaves per species and sex were photographed (Nikon AF-S 60mm f/2.8 ED Micro) with a scale bar, and surface area measured in ImageJ (Schneider et al. 2012). Leaves were then dried and weighed on an analytical balance (Mettler Toledo XP2003SDR). Leaf dimorphism was regressed against the number of male-biased and female-biased transcripts using a phylogenetic least squares model (gls), fitted by maximum-likelihood in the R package nlme (Pinheiro et al. 2018) and using the species tree inferred in the present study.

#### Reconstruction of ancestral states

To infer where along the species tree sex-biased expression had evolved, we reconstructed the ancestral states for each sex-biased gene, with the three discrete character states ‘unbiased’, ‘male-bias’ and ‘female-bias’, using maximum-likelihood implemented in the ace function in the R package APE v5.3 (Paradis and Schliep 2019).

#### Permutation tests for number of shared sex-biased genes

We devised a permutation procedure to test whether the observed frequencies and gene identities of repeated and divergent evolution of sex-bias were different from a random expectation. Cases in which two or more species share sex bias for a gene, and if this sex bias is not due to ancestral sex bias, can be interpreted as repeated evolution and suggest sex-related functional relevance. However, if the identities (and hence putative functions) of sex-biased genes were irrelevant, we may nevertheless expect some coincidental overlap in sex-biased genes between species. This random expectation is analogous for genes with divergent sex-bias. We generated such random expectations by permuting the identities of sex-biased genes within each species 10,000 times while keeping the numbers intact, and scoring the interspecific overlaps. Four genes with an ancestral gain of sex-biased expression were excluded from this analysis. P-values were quantified as the proportion of overlaps in permuted datasets that were greater or equal to the observed overlaps.

#### Functional gene annotation

Majority-consensus peptide sequences were generated for the PREQUAL-cleaned and MAFFT-aligned orthogroups. These were annotated against *Arabidopsis thaliana* genes (Araport11.201606, www.arabidopsis.org) (Cheng et al. 2017) using BLASTP, retaining only best hits with an e-value threshold of 10^-5^. A broad level functional categorization of the SBGs annotated to *A. thaliana* genes was generated through the TAIR web portal https://www.arabidopsis.org/tools/bulk/go/index.jsp, and the results were visualised in ComplexHeatmap. We also retrieved functional annotation based on the *Arabidopsis* gene identifiers using DAVID (Huang et al. 2009), and clustered the default annotations at medium stringency.

#### Phylogenetically independent contrasts

The mean of the absolute standardized phylogenetically independent contrasts (PICs, (Felsenstein 1985)) per gene was employed as a measure of the rate of expression evolution (Garland 1992).

For each gene, PICs were calculated based on the species tree and the mean log_2_(TMM-TPM + 1) expression values per species, using the pic function in APE. Sex-biased expressions themselves were excluded when calculating PICs, so that the PICs only measure gene expression variation without sex-bias. Significance of the mean difference between SBGs and unbiased genes in the mean absolute PICs was assessed on the basis of 10,000 permutations, using the function permTS in the R package perm v1.0 (Fay and Shaw 2010).

#### Expression distances as a function of sequence divergence

Gene expression distances between pairs of species were calculated as 1 – Pearson’s correlation coefficient, using the mean log_2_(TMM-TPM + 1) expression levels per species and sex. This was done for three different categories, namely for unbiased genes with the average expression levels over both sexes, for SBGs with the male expression levels, and for SBGs with the female expression levels. Note that for SBGs, this includes only expression levels and species in which these genes were not sex-biased; hence sex-biased expression itself did not contribute to the gene expression distance for SBGs. For visualisation of correlation trends, linear models were fitted for expression distances as a function of sequence divergence using ggplot2 (Wickham et al. 2020), or the lm function in R. Significance of the correlations between pairwise distance matrices was evaluated by Mantel tests in vegan v2.5-6 (Oksanen et al. 2019) with 100,000 permutations.

We conducted a number of tests to verify that the higher expression evolutionary rates observed for SBGs were not spurious artefacts. First, we investigated correlations between inter-group and intra-group variation in gene expression, using simulated expression levels and real expression data. Second, we compared the mean coefficient of variation (CV) of expression counts between SBGs and unbiased genes within each species and sex using the function permTS in the R package perm, on the basis of 10,000 permutations. Third, we tested whether greater expression noise alone could explain the differences between sex-biased and unbiased genes in the linear model coefficients for gene expression distance between species as a function of sequence divergence between species. Hence, we compared the observed linear model intercept and slope estimates with artificial estimates from 1,000 datasets in which the actual sex-biased genes were replaced by an equal number of randomly chosen unbiased genes that matched the expression noise level of the true sex-biased genes (i.e., in each artificial dataset, for each sex-biased gene, one unbiased genes was sampled that matched the sex-biased gene within 95% – 105% of its coefficient of variation in expression over all species and sexes). Fourth, the empirical falsepositive rate for SBGs was determined by conducting the DGE analyses between the sexes (as described above) 1,000 times, with datasets in which the sex was randomised (re-sampling males and females without replacement).

#### Putative adaptive shifts in expression level

As an indicator of directional selection on gene expression level, we used the delta-x statistic of (Zemp et al. 2016), which here is the ratio of the absolute difference in mean expression between two groups (divergence) over the standard deviation of expression within a focal group (diversity / polymorphism). The rationale is analogous to indicators of selection on DNA sequence variation (McDonald and Kreitman 1991), i.e., directional selection on a trait (expression level) should both increase between-group divergence and decrease within-group variation, whereas traits evolving mainly under drift should show variation similar or greater than divergence. Delta-x was calculated on the basis of log_2_(TMM-TPM + 1) expression levels, separately for each gene, each *Leucadendron* species, and for males and females. Divergence was estimated as the difference between the mean expression levels in the sister species (as defined on the species tree), and diversity as the standard deviation over the six (or five) replicates of the focal species and sex. We classified divergences as quantitative evolutionary shifts if the interspecific fold-change of the means was at least 1.5. Shifts in expression showing a delta-x greater than 5.0 were classified as ‘high delta-x’, i.e., a category more consistent with adaptive evolution. Counts of sex-biased shifts and unbiased shifts in the two categories ‘high delta-x’ and ‘low delta-x’ were compared using a 2 x 2 chi-squared test with Yates’s continuity correction in R.

#### Expression specificity

The expression specificity for *Leucadendron* genes over different tissues and developmental stages was transferred from *Arabidopsis thaliana* genes, as calculated over 24 different groups of tissues and developmental stages in the gene expression atlas of (Klepikova et al. 2016). The original Shannon entropy values were transformed to a scale ranging from 0 (ubiquitous expression) to 1 (tissue- or stage-specific expression), as proposed by (Kryuchkova-Mostacci and Robinson-Rechavi 2017). The difference in mean expression specificity between SBGs and unbiased genes was assessed by 10,000 permutations using the function permTS in the R package perm.

#### Rates of coding sequence evolution

We selected orthogroup gene trees and clean alignments (see above section ‘Species tree and interspecific sequence divergence’) with at least four represented species. Omega (dN/dS) was estimated with PAML (Yang 2007) using branch model 2 as implemented in the python package ete3 (Huerta-Cepas et al. 2016), on alignments of one foreground sequence and as background sequences the three closest orthologs in the orthogroup. A single omega value per species and orthogroup was reported as the mean omega of all sequences of a species per orthogroup. Thus, our estimate of omega of an orthogroup in a focal species describes how the sequences of that orthogroup, on average, differ from the closest sequences in different species, i.e., any evolution leading towards the focal species. Our estimate is therefore comparable to the omega from PAML’s yn00 model, which is widely reported in studies of two species with one-to-one orthology, although we analysed multiple species with potentially multiple sequences each. We tested for differences in mean omega between unbiased, male-biased and female-biased cases on the basis of 1,000 permutations using the function permTS in the R package perm.

**Fig. S1.**
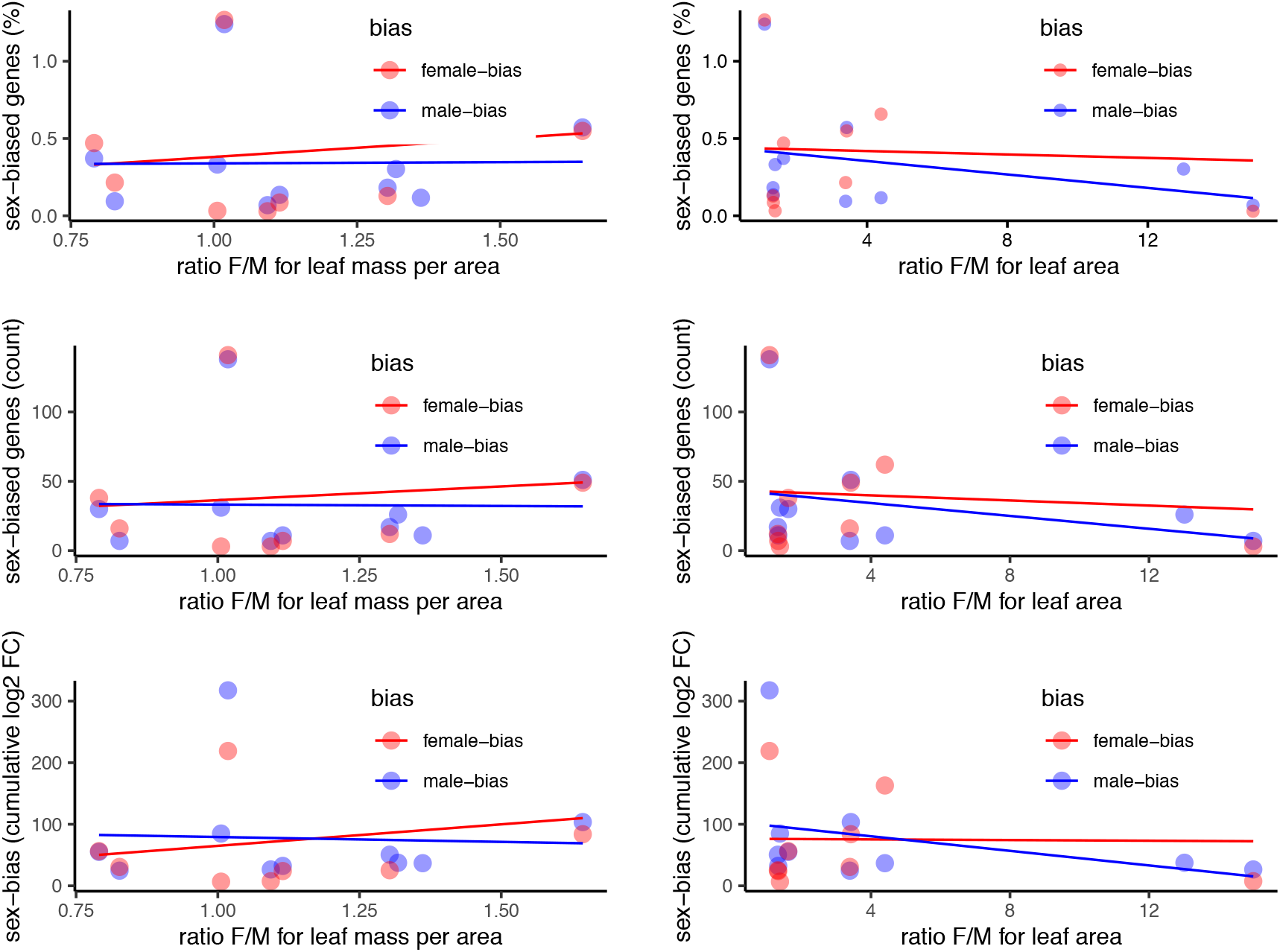
All six alternative scatter plots for sex-biased gene expression as a function of morphological sexual dimorphism in the leaves of ten species of *Leucadendron*, complementing Figure 1 C) of the main text. The underlying data is found in Table S3.

**Fig. S2.**
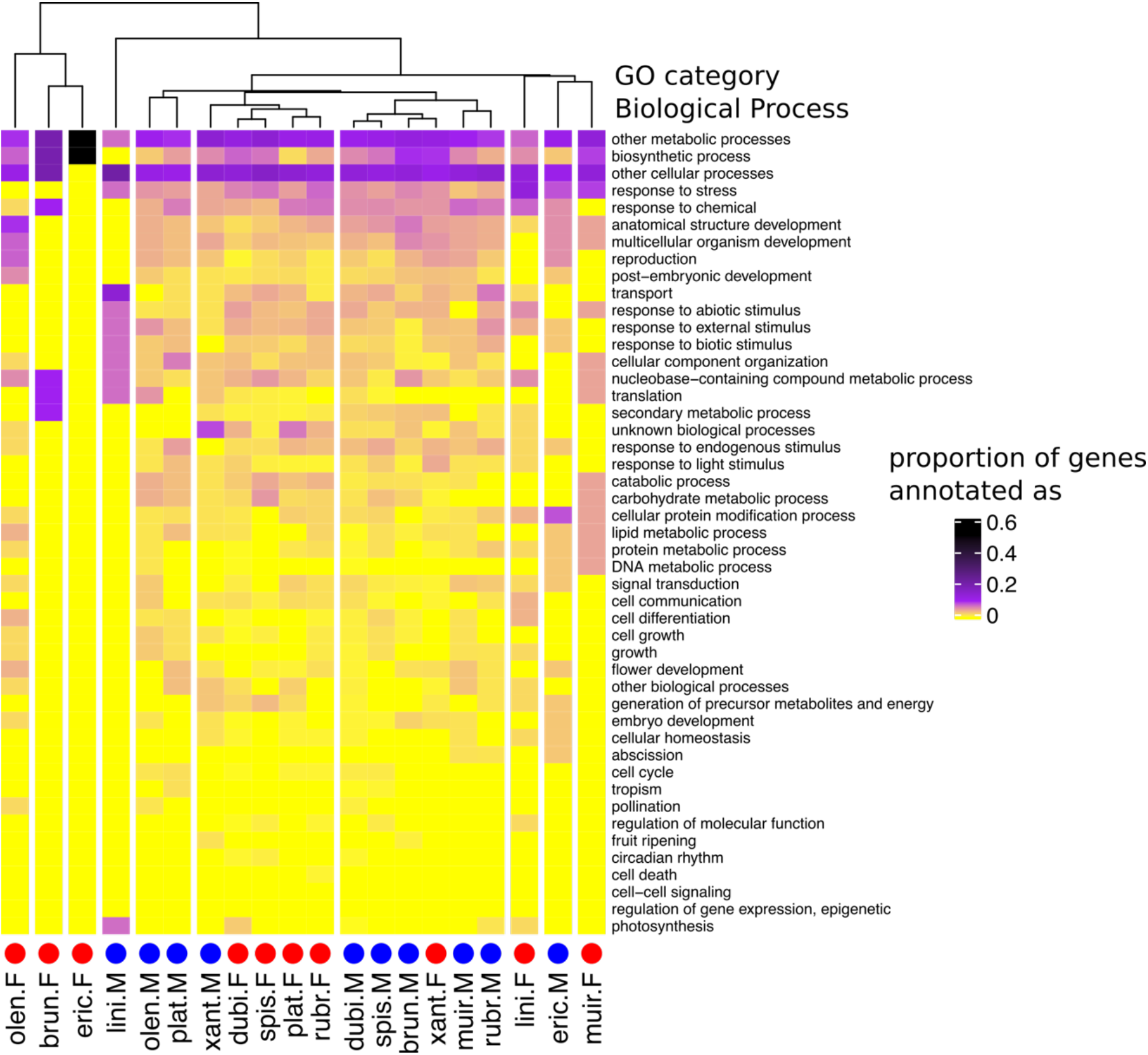
Heatmap and cluster dendrogram for biological processes putatively regulated by sex-biased genes in ten *Leucadendron* species. Values are the proportions of sex-biased genes annotated with a given biological process (functional categorization according to TAIR database). The cluster dendrogram is based on Pearson’s correlation coefficient.

**Fig. S3.**
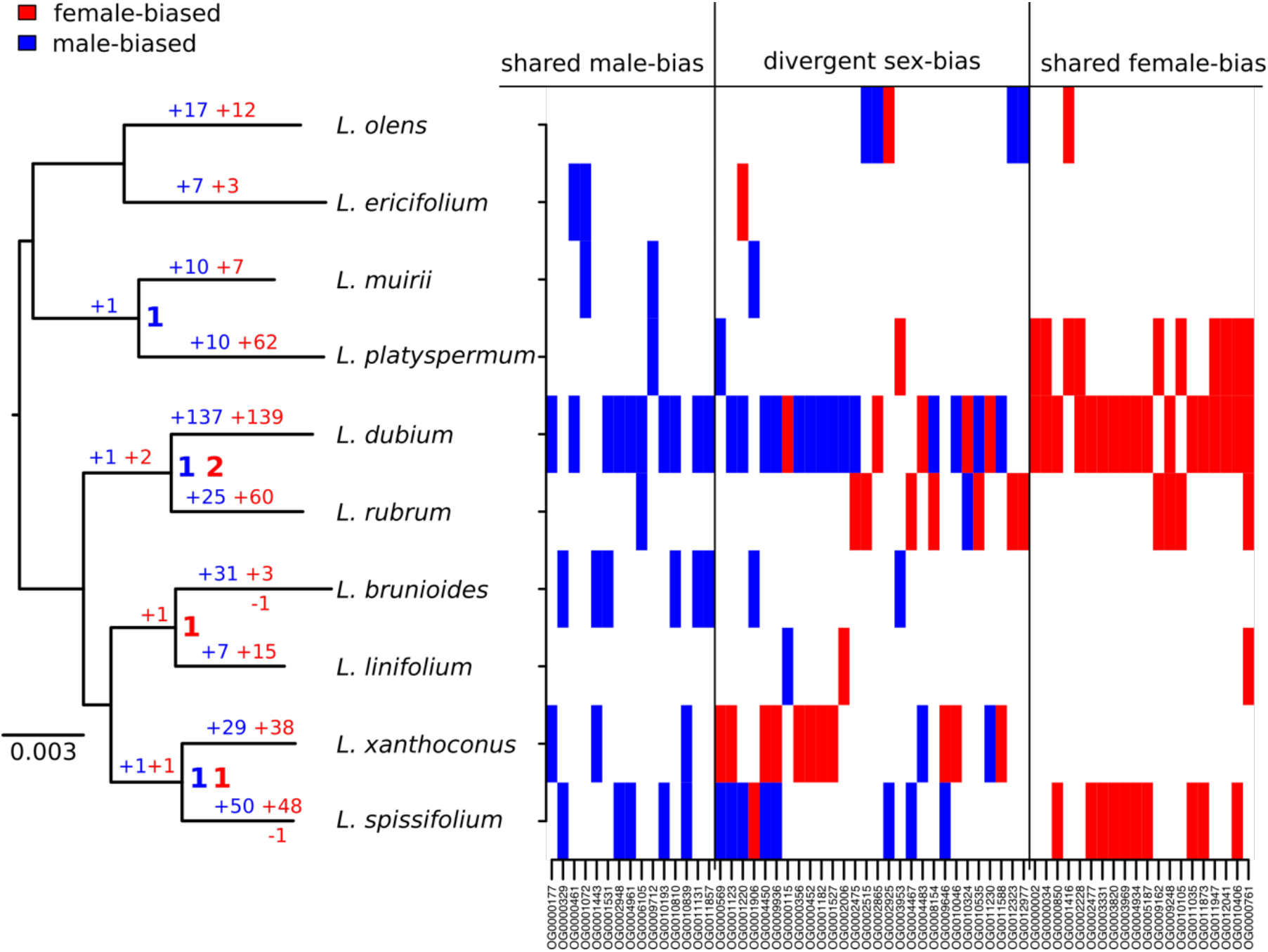
Summary of evolutionary histories inferred for SBGE in *Leucadendron*. Left: species tree annotated with inferred numbers of sex-biased genes at ancestral nodes (bold), and gains of sex-biased genes annotated on each branch; no losses were inferred. Right: Table marking sex-biased status for the 63 genes which showed sex-bias in more than one species, either shared bias in the same direction, or divergent sex-bias. More details can be found in Table S4.

**Fig. S4.**
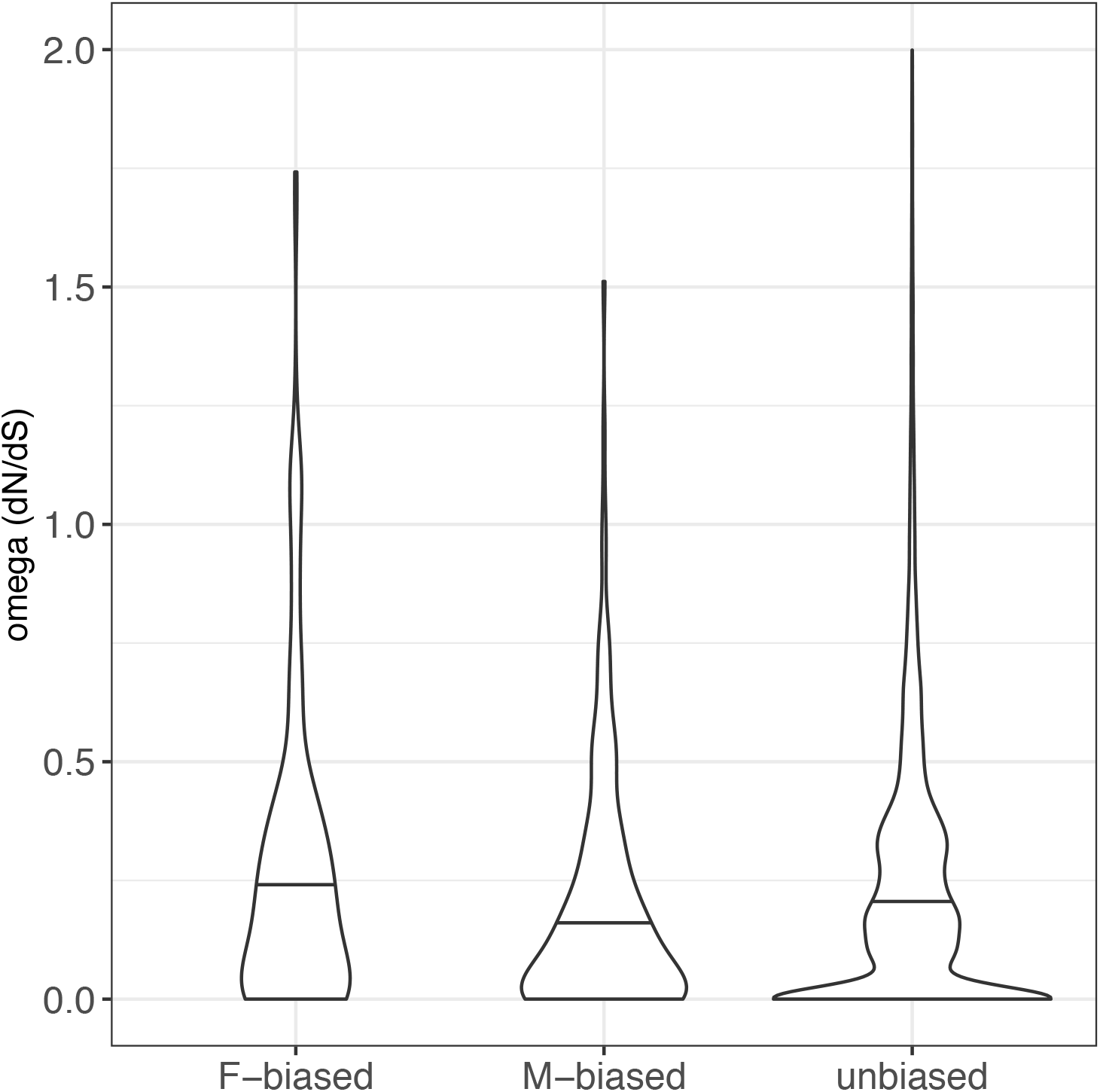
Molecular sequence evolution measured as dN/dS (omega) for ten *Leucadendron* species and all expressed genes in sex-biased and unbiased categories (horizontal bar in the violin is the median). Mean omega was not significantly different between unbiased and sex-biased genes (permutation tests p > 0.05).

**Fig. S5.**
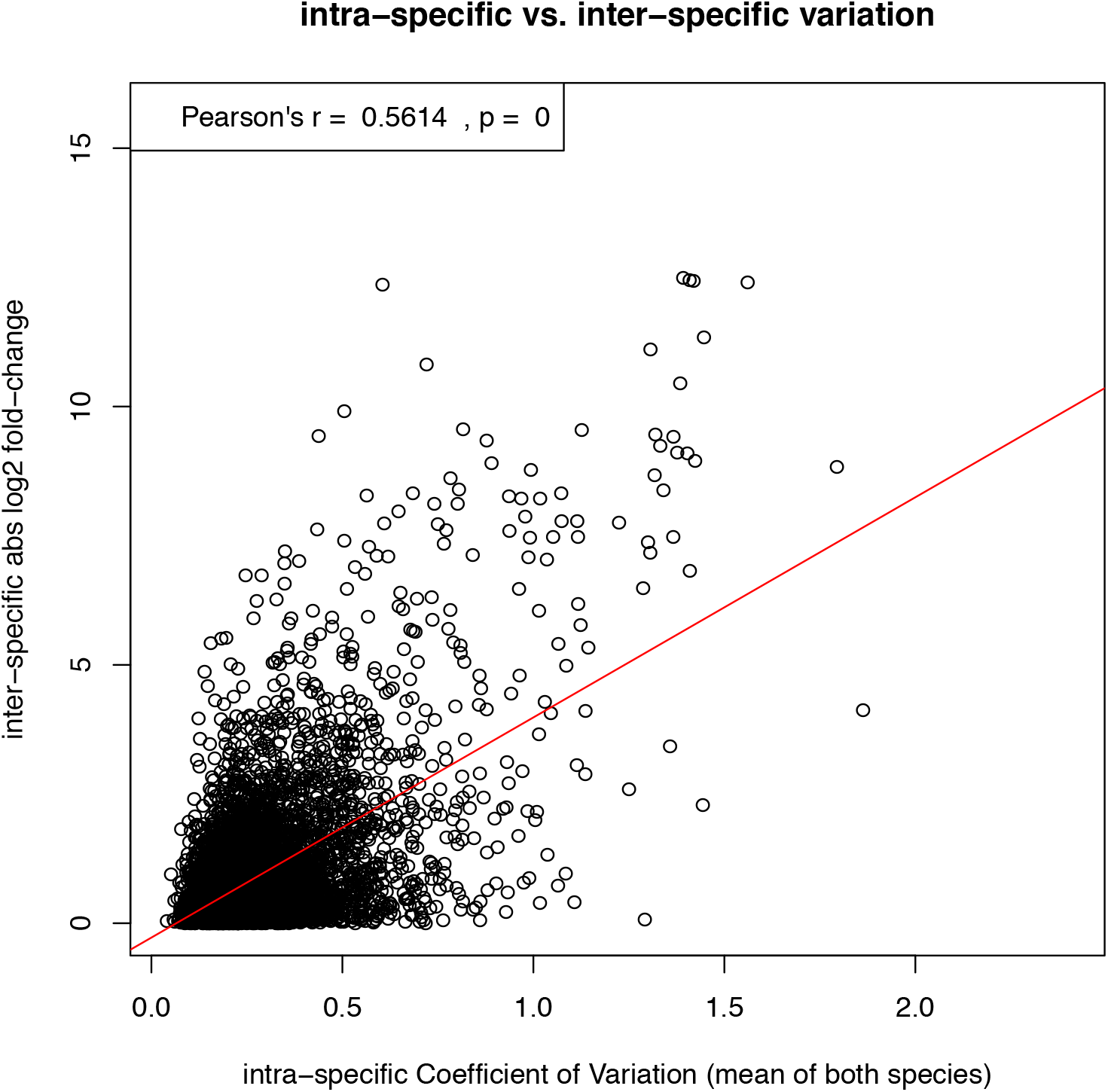
Demonstration that inter-specific differences in expression level tend to correlate positively with the level of intra-specific expression variation, posing the risk that genes with noisy expression are mistakenly inferred to have fast rates of expression evolution. This example scatterplot and correlation test shows all expressed genes in the species pair *L. rubrum* and *L. dubium*. The X-axis shows the intra-specific Coefficient of Variation over all 12 replicates per species, averaged over both species. The Y-axis shows the inter-specific log_2_ fold-change between the mean expression levels of the two species. Similar results were obtained for all other pairs of species and also with entirely artificial, randomly simulated data (not shown). However, as described in the main text and Figure S6, this trend did not confound the inference of faster expression evolution for sex-biased genes in this study, because sex-biased genes showed low expression noise (i.e. expression variation within sexes).

**Fig. S6. (separate file Figure_S6.pdf).** Comparisons of Coefficient of Variation per sex for sex-biased and unbiased genes, separately for each *Leucadendron* species.

**Fig. S7. (separate file Figure_S7.pdf).** Comparisons of linear model coefficients for gene expression distance between species as a function of sequence divergence between species, for sex-biased genes and unbiased genes. Shown are the observed intercept and slope estimates, together with histograms of the same estimates for 1,000 datasets in which the actual sex-biased genes were replaced by an equal number of unbiased genes that are matched to the expression noise level of sex-biased genes (i.e. within 95-105% of their coefficient of variation in expression over all species and sexes).

**Fig. S8.**
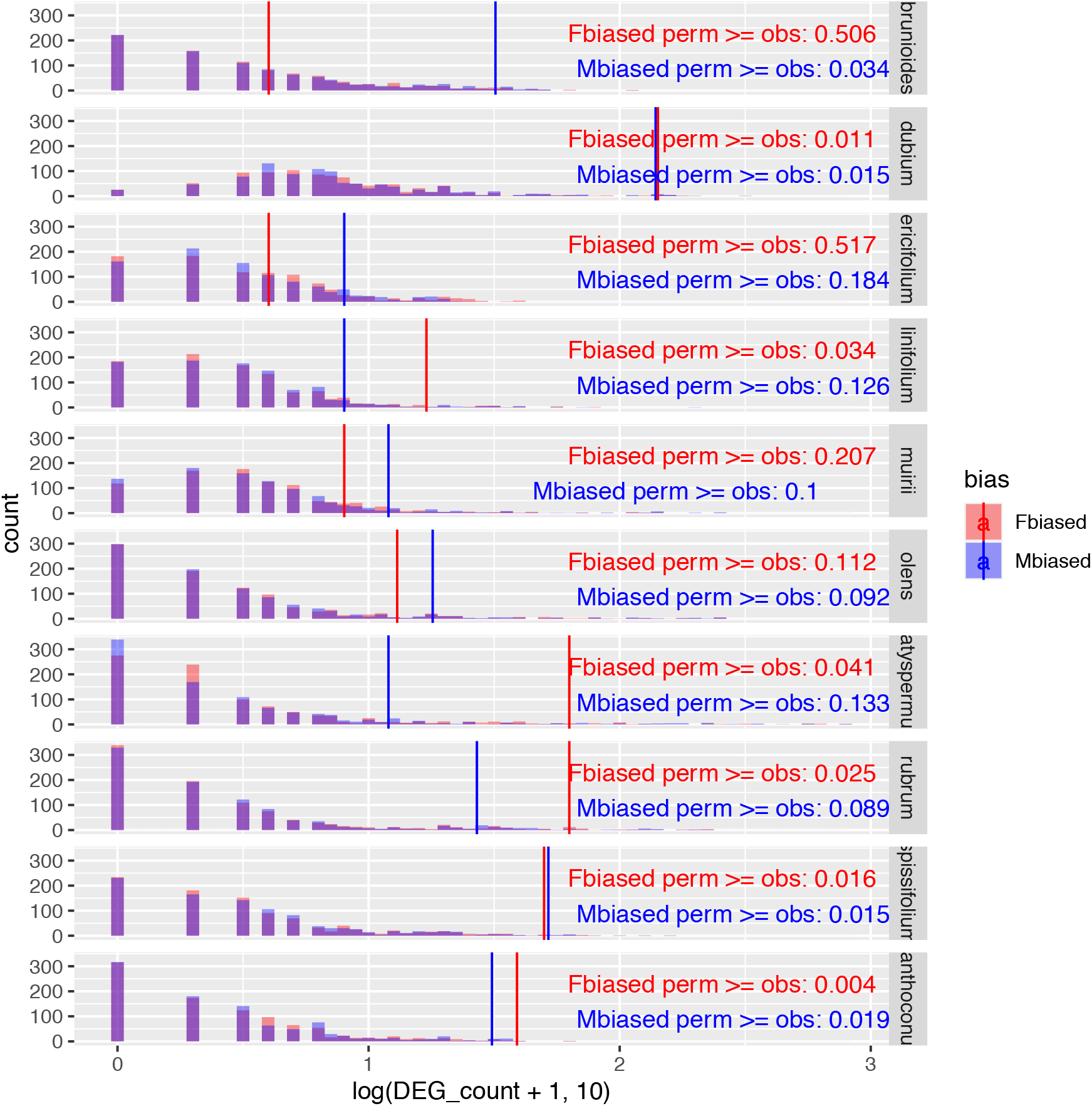
Challenging the inference of sex-biased genes by repeated permutation of the sexes. In most species and sexes, observed counts of sex-biased genes are significantly greater than counts from permuted datasets, indicating that the observed sex-biased genes are unlikely to represent stochastic artefacts.

**Table S1 (separate file).** Descriptive statistics of denovo assembled transcriptome sequences.

**Table S2 (separate file).** List of all 650 sex-biased genes and direction of sex-bias per *Leucadendron* species, together with annotated AT gene identifier.

**Table S3 (separate file).** Percentages, counts and cumulative fold-changes of sex-biased genes, and metrics of morphological leaf dimorphism for each of the *Leucadendron* species.

**Table S4 (separate file).** Overview of the 63 genes that were sex-biased in two or more species among the set of ten *Leucadendron* species.

**Table S5 (separate file).** Functional annotations and annotation clusters using the default annotations at medium stringency in the DAVID database for male-biased genes in *Leucadendron*.

**Table S6 (separate file).** Functional annotations and annotation clusters using the default annotations at medium stringency in the DAVID database for female-biased genes in *Leucadendron*.

**Table S7 (separate file)**. Functional annotations and annotation clusters using the default annotations at medium stringency in the DAVID database for for sex-biased genes per each of ten *Leucadendron* species.

